# Causal evidence for a double dissociation between object- and scene-selective regions of visual cortex: A pre-registered TMS replication study

**DOI:** 10.1101/2020.07.24.219386

**Authors:** Miles Wischnewski, Marius V. Peelen

## Abstract

Natural scenes are characterized by individual objects as well as by global scene properties such as spatial layout. Functional neuroimaging research has shown that this distinction between object and scene processing is one of the main organizing principles of human high-level visual cortex. For example, object-selective regions, including the lateral occipital complex (LOC), were shown to represent object content (but not scene layout), while scene-selective regions, including the occipital place area (OPA), were shown to represent scene layout (but not object content). Causal evidence for a double dissociation between LOC and OPA in representing objects and scenes is currently limited, however. One TMS experiment, conducted in a relatively small sample (N=13), reported an interaction between LOC and OPA stimulation and object and scene recognition performance (Dilks et al., 2013). Here, we present a high-powered pre-registered replication of this study (N=72, including male and female human participants), using group-average fMRI coordinates to target LOC and OPA. Results revealed unambiguous evidence for a double dissociation between LOC and OPA: Relative to vertex stimulation, TMS over LOC selectively impaired the recognition of objects, while TMS over OPA selectively impaired the recognition of scenes. Furthermore, we found that these effects were stable over time and consistent across individual objects and scenes. These results show that LOC and OPA can be reliably and selectively targeted with TMS, even when defined based on group-average fMRI coordinates. More generally, they support the distinction between object and scene processing as an organizing principle of human high-level visual cortex.

**Significance Statement:** Our daily-life environments are characterized both by individual objects and by global scene properties. The distinction between object and scene processing features prominently in visual cognitive neuroscience, with fMRI studies showing that this distinction is one of the main organizing principles of human high-level visual cortex. However, causal evidence for the selective involvement of object- and scene-selective regions in processing their preferred category is less conclusive. Here, testing a large sample (N=72) using an established paradigm and a pre-registered protocol, we found that TMS over object-selective cortex (LOC) selectively impaired object recognition while TMS over scene-selective cortex (OPA) selectively impaired scene recognition. These results provide conclusive causal evidence for the distinction between object and scene processing in human visual cortex.

## Introduction

Our daily-life environments, such as city streets, forests, and kitchens, are characterized both by local object content (cars, trees, refrigerators) and global spatial layout. These sources of information are to some extent independent, with many scenes being identifiable without requiring individual object recognition, and most objects being identifiable without a background scene. Accordingly, cognitive neuroscience research has revealed distinct pathways in visual cortex that are primarily involved in either scene or object recognition (Oliva, 2013; Epstein, 2014).

In humans, fMRI research has shown that viewing objects (vs scrambled controls) selectively activates regions in lateral occipital cortex (LOC) and posterior fusiform gyrus (Malach et al., 1995; Grill-Spector, 2003). By contrast, viewing scenes (vs objects) activates three scene-selective regions (Epstein and Baker, 2019): the parahippocampal place area (PPA), the retrosplenial complex (RSC) and the occipital place area (OPA). Studies using multivariate pattern analysis have shown that these scene- and object-selective regions represent distinct aspects of a scene, with scene-selective regions representing spatial layout (e.g., open vs closed) and object-selective regions representing object properties (Park et al., 2011; Kravitz et al., 2011; Harel et al., 2013).

While the neuroimaging evidence for a functional distinction between scene- and object-selective regions is convincing, the evidence for these regions being causally involved in selectively processing their preferred category is weaker. A number of studies have used transcranial magnetic stimulation (TMS) to interfere with activity in one of these regions to demonstrate impaired object or scene recognition. These studies showed that TMS over LOC impaired object recognition performance (Pitcher et al., 2009; Mullin and Steeves, 2011; Koivisto et al., 2011). Due to physical limitations of TMS reaching deeper structures, studies investigating scene recognition have targeted OPA, rather than PPA or RSC. These studies found that scene categorization is impaired when OPA is stimulated (Dilks et al., 2013; Ganaden et al., 2013). The causal evidence for a double dissociation between object-selective LOC and scene-selective OPA, however, is limited to a single experiment, conducted in a relatively small sample (N=13; Dilks et al., 2013). Providing such evidence is important for ruling out alternative explanations for impaired performance. For example, TMS-related artifacts (e.g., discomfort) differ depending on scalp location, compromising the direct comparison of task performance across regions. TMS-related artifacts (e.g., the clicking sound of a TMS pulse) may also differentially affect different tasks, for example related to the attentional requirements of each task.

In the experiment by Dilks et al. (2013), participants performed two four-alternative forced-choice (4AFC) categorization tasks, one involving scenes (beach, forest, city, kitchen) and one involving objects (camera, chair, car, shoes). In different blocks, TMS was applied over right LOC, right OPA, or vertex. Results showed a significant interaction between region (LOC, OPA, vertex) and task (scenes, objects). However, such an interaction does not necessarily indicate a double dissociation as it could, in principle, result from task-specific effects of TMS over one region alone. Furthermore, TMS over LOC has been shown to both impair object perception and facilitate scene perception (Mullin and Steeves, 2011), such that scene recognition might be worse when stimulating OPA as compared to LOC even when TMS over OPA has no effect on scene recognition. Comparisons with vertex (control) stimulation are therefore important for interpreting the interaction. These comparisons provided some evidence that TMS over LOC impaired object recognition (*p* = 0.06) and TMS over OPA impaired scene recognition (*p* < 0.05).

While these results are encouraging, effects of TMS are variable across individuals (Ridding & Zieman, 2010) and effect sizes are typically small, such that large samples are needed to obtain generalizable results. Indeed, it is increasingly recognized that results obtained in small samples are often not reproducible or overestimate effect sizes (Button et al., 2013). Therefore, to provide conclusive causal evidence for a double dissociation between object- and scene-selective regions, we replicated the experiment by Dilks et al. (2013) in a large sample of participants (N=72) using a pre-registered protocol.

## Materials and Methods

The pre-registration is available at this website: https://aspredicted.org/jm6ey.pdf.

### Participants

Seventy-two volunteers (mean age± SEM = 23.30 ± 0.42, range 18-33 years, 43 female) participated in the study. The current study was performed to select participants for a subsequent study, for which three groups of 24 participants were needed. All volunteers had normal or corrected-to-normal vision and were right handed. Exclusion criteria were: (i) History of epileptic seizures, (ii) a nuclear family member afflicted with epilepsy, (iii) history of psychiatric or neurological diseases, (iv) metallic objects in the head, (v) implanted devices (e.g., pacemaker, cochlear implant), and (vi) having used psycho-active medication or recreational drugs less than 48 h before the experiment. The protocol was approved by the medical ethical committee of the Commissie Mensgebonden Onderzoek, Arnhem-Nijmegen and carried out in accordance with the Declaration of Helsinki (Fortaleza amendments).

### Behavioral task

Participants performed a 4AFC task, identical to the task used by Dilks et al. (2013). In a blocked design, participants were either presented with an object, which could be a camera, car, chair or shoes, or with a scene, which could be a beach, city, forest, or kitchen. For all stimuli, grayscale images from the SUN Database (Xiao et al., 2010) were used, which were further degraded by blending the image with 8×8 grid of tiles. For scenes, these tiles were grayscale with random intensity. For objects, a scrambled version at 60% transparency was used as tiles and overlaid on the image. All images spanned a visual angle of 9×9 degrees.

The experiment consisted of two parts. The first part of the experiment determined the optimal stimulus presentation duration by using a thresholding procedure that leads to an average score of ~63% for each category. For this a QUEST staircase procedure was used (number of trials = 32; beta = 3.5; delta = 0.01; gamma = 0.5; grain = 2) in two runs of 256 stimuli (i.e. 128 stimuli for objects as well as for scenes, equaling 32 stimuli per category). The range of stimulus presentation was fixed between 64-128 ms per category. The lower (shorter presentation time) threshold value of the two runs for each category was selected for the second part of the experiment. The stimulus presentation times on average were: camera 91.6 ms, car 74.9 ms, chair 114.0 ms, shoes 110.5 ms, beach 80.8 ms, city 93.3 ms, forest 86.9 ms, and kitchen 116.1 ms. Each stimulus was followed by a mask, which was presented for 500 ms. The mask was made of a 4×4 grid of tiles containing a scramble of 8 randomly selected image parts. For scenes, undegraded images were used, whereas for objects, degraded images were used to make the mask.

The second experimental part consisted of six runs of 64 stimuli (i.e. 32 stimuli for objects as well as for scenes, equaling 8 stimuli per category). During this part participants were stimulated with TMS, which started at stimulus onset. Stimulation site, that is, rLOC, rOPA, or vertex, was changed each run and followed a pseudorandom, counterbalanced palindromic design.

### Transcranial magnetic stimulation and site localization

A MagVenture MagPro X100 was used to deliver TMS with a Cool B-65 figure-of-eight coil. Note that this is different from Dilks et al. (2013), who used a MagStim Super Rapid^2^ stimulator.

At the onset of each object or scene stimulus a train of 5 pulses was applied at 10 Hz (i.e., 500 ms total duration of a single train) at 60% of maximum stimulator output (MSO). TMS was delivered over rLOC, rOPA and vertex, with a posterior-to-anterior magnetic field orientation [10]. The correct location during stimulation was monitored using a Localite neuronavigation system. The size and shape of a participant’s head was modelled by marking left and right tragus of the ear, left and right canthi of the eyes, nasal bridge, inion and vertex. Subsequently, a standard MNI brain was fit, based on those points. MRI coordinates were used to identify the peak activation voxel of the rLOC, that is Talairach 45,-74,0 (Pitcher et al., 2009), and of the rOPA, that is Talairach 34,-77,21 (Julian et al., 2016). Note that Dilks et al. (2013) used individual fMRI coordinates. The location of the vertex was determined as the midpoint between both tragi, on top of the head. For all stimulation sites, coil location was monitored using the Localite system and was kept constant within a range of 2 mm displacement.

Before the start of the task, the phosphene threshold (PT) of each participant was determined using the ‘method of constant stimuli’ (Mazzi et al., 2017). On average, PT was 60.43 ± 1.45% of MSO ranging between 38 and 100 MSO. After PT determination the 5 pulse, 10 Hz stimulation, as during the experiment, was shortly applied to each experimental region for participants to acclimatize to the feeling of TMS. No adverse events were reported.

### Statistical analysis

Similar to Dilks and colleagues (2013), for the main analysis a 3×2 GLM repeated measures ANOVA was performed with percentage correct as dependent variable, following our pre-registration. TMS site (rLOC, rOPA and vertex) and category (object and scene) acted as independent variable. A significant main and interaction effects were followed by post hoc t-tests. In addition, Bayes factors (BF_10_) are reported for post hoc analysis. Values larger than 1 indicate evidence that the alternative hypothesis (i.e., performance in the active TMS condition is different from vertex TMS) is true, with larger values suggesting stronger evidence. Values smaller than 1 indicate evidence for the null hypothesis (i.e., performance does not differ between active and vertex TMS) with values closer to 0 suggesting stronger evidence. Data was further explored by comparing the LOC effect (percentage correct during vertex – LOC stimulation during object recognition) and OPA effect (percentage correct during vertex – OPA stimulation during scene recognition) between first and second half of the experiment. A paired samples t-test was used to test for differences between halves and one sample t-tests with zero as test value was used to investigate whether the effect was larger than zero. Finally, repeated measures ANOVAs were used to test for differences between categories for the LOC and OPA effect as well as a one-sample t-test to indicated whether the effect is larger than zero. All analyses were performed using SPSS 25.0 (ANOVA and post hoc tests) and JASP 0.12.2 (Bayes factors).

## Results

### Main results: Replication of Dilks et al (2013)

A 3 (TMS site: rOPA, rLOC, vertex) x 2 (Category: objects, scenes) ANOVA revealed a significant interaction (F(2,142) = 7.09, p = 0.001), indicating that TMS site differentially affected performance (% correct) in the two tasks. To test whether object and scene recognition performance was impaired following stimulation, we followed up this interaction by separate one-way ANOVAs for each category.

For the object recognition task, a significant main effect of TMS site indicated that object recognition was affected by TMS (F(2,142) = 8.38, p < 0.001, Figure 2A). Post hoc t-tests showed that performance was reduced during LOC TMS compared to vertex TMS (t(71) = 4.05, p < 0.001, d = 0.48, BF_10_ = 269.65, Figure 2B) and compared to OPA TMS (t(71) = 2.55, p = 0.013, d = 0.30, BF_10_ = 3.81). OPA TMS did not impair object recognition performance compared to vertex TMS (t(71) = 1.42, p = 0.159, d = 0.17, BF_10_ = 0.31).

**Figure 1.**
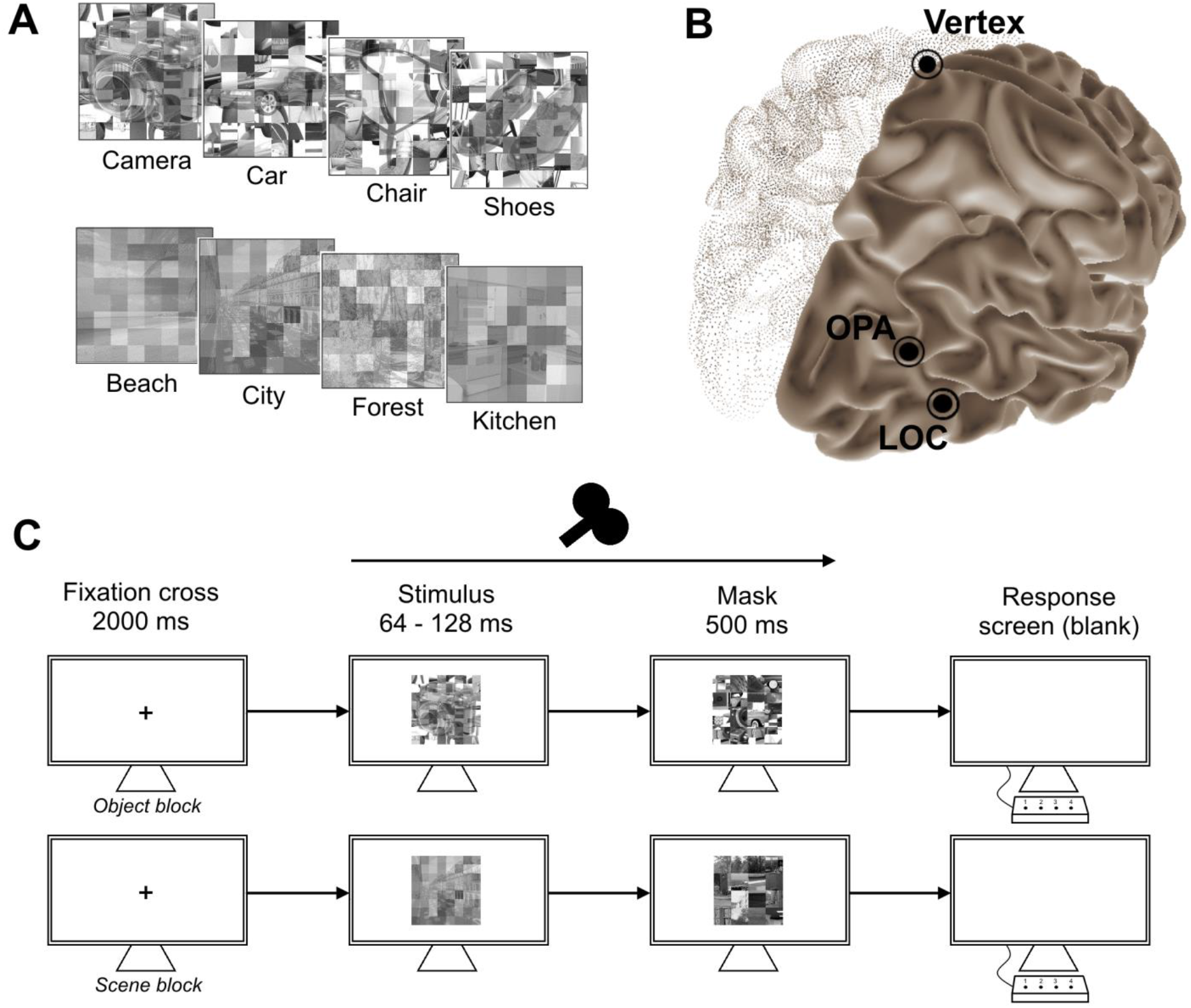
A) Examples of stimuli for each category. B) TMS sites used in the present study. OPA and LOC were based on fMRI coordinates from previous studies, i.e. for the LOC Talairach 45,-74,0 (Pitcher et al., 2009) and for the OPA Talairach 34,-77,21 (Julian et al., 2016). C) Example trial of the 4AFC object/scene recognition task. Five TMS pulses at a rate of 10 Hz were delivered at the onset of the stimulus.

**Figure 2.**
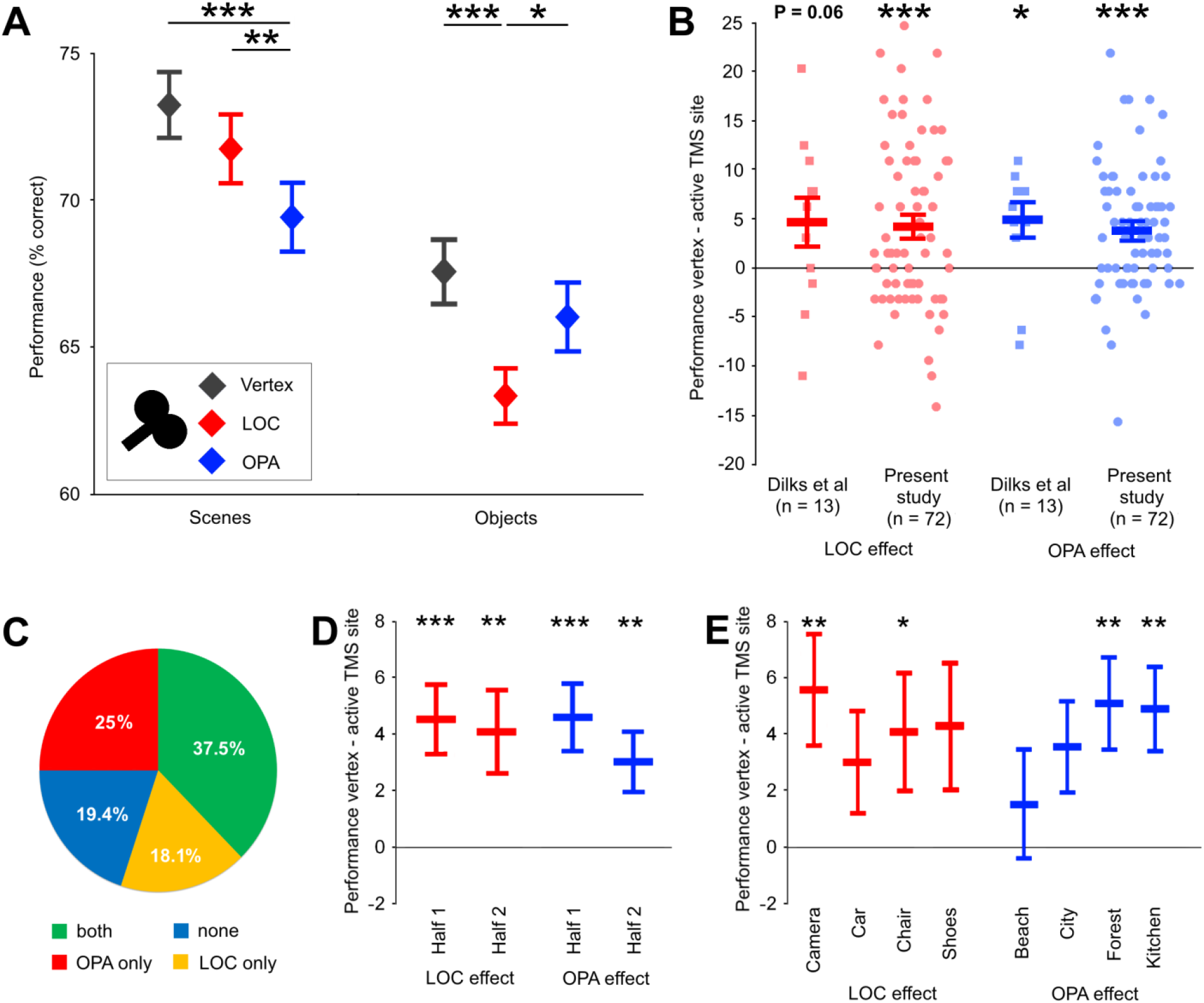
A) Results of the present study (N = 72). The findings are similar to Dilks et al. (2013) showing decreased performance for scene recognition during TMS over OPA (compared to vertex) and decreased performance for object recognition during TMS over LOC (compared to vertex). B) Average and individual data of the LOC effect (% performance difference vertex – LOC during object recognition) and OPA effect (% performance difference vertex – OPA during scene recognition) for both the present study and Dilks et al. (2013). C) Percentages of participants susceptible to TMS over OPA, LOC, both or neither. D) LOC (red) and OPA (blue) effect for the first half and second half of the experiment. E) LOC (red) and OPA (blue) effect per category. In all graphs: * p < .05, ** p < .01, *** p < .001, with error bars reflecting the SEM.

For the scene recognition task, we similarly found a main effect of TMS site (F(2,142) = 11.97, p < 0.001, Figure 2A). In this case, performance was reduced during OPA TMS compared to vertex TMS (t(71) = 4.91, p < 0.001, d = 0.58, BF_10_ = 5753.48, Figure 2B) and compared to LOC TMS (t(71) = 3.08, p = 0.003, d = 0.36, BF10 = 5.39). LOC TMS did not impair scene recognition performance compared to vertex TMS (t(71) = 1.51, p = 0.135, d = 0.18, BF_10_ = 0.687).

These results replicate the results of Dilks et al. (2013), providing convincing causal evidence for a double dissociation between scene- and object-selective regions in the recognition of scenes and objects. The overall pattern of results of the two studies was nearly identical, with statistically stronger evidence in the current study due to the larger sample size (Figure 2B).

### Individual differences and consistency across time

Although overall TMS was effective in altering scene and object recognition performance, there is a notable amount of individual variability in response to TMS. To get an idea of this variability, the responder rate was calculated. 37.5% of participants showed both an effect of TMS over LOC (object recognition performance vertex – LOC > 0) and OPA (scene recognition performance vertex – OPA > 0), 18.1% of participants showed an effect of LOC but not OPA TMS, and 25.0% of participants showed an effect of OPA but not LOC TMS (Figure 2C). This means that the susceptibility to TMS over LOC and OPA was 55.6% and 62.5%, respectively. Although these rates are lower than those of Dilks and colleagues (69.2% and 84.6%, respectively), chi-square tests indicated no significant difference between these percentages (X^2^ = 0.84, p = 0.358, and X^2^ = 2.40, p = 0.122, respectively). The susceptibility rates found here are in agreement with susceptibility rates of previous studies targeting visual areas, such as Van Koningsbruggen et al. (2013), who showed a susceptibility rate of 54% for TMS over the extrastriate body area (EBA), located near the LOC.

Individual differences in the excitability of the stimulated cortex, skull thickness, neuronal and axonal orientation as well as metabolic factors may impact the susceptibility to TMS. One way to control for this variability is to take into account the phosphene threshold. We found that phosphene threshold was not significantly correlated with the LOC TMS effect (r = 0.04, p = 0.739), nor with the OPA TMS effect on performance (r = 0.13, p = 0.269). This suggests that the above-mentioned physiological differences between participants, to the extent that these are captured by phosphene threshold, did not significantly mediate our effects.

Since all TMS sites were stimulated twice in a palindromic design it was possible to investigate the consistency of the effects over time by comparing the TMS-induced LOC effect (% correct object recognition, vertex – LOC) and OPA effect (% correct scene recognition, vertex – OPA) in the first half to the second half of the task. The LOC effect was significant for both the first (4.60 ± 1.15%, t_0_(71) = 3.90, p < 0.001, d = 0.47) and second half (4.04 ± 1.40%, t_0_(71) = 2.88, p = 0.005, d = 0.34), with no significant difference between the two halves (t(71) = 0.36, p = 0.724, d = 0.04, BF_10_ = 0.138, Figure 2D). Similarly, the OPA effect was significant in the first (4.69 ± 1.12%, t0(71) = 4.18, p < 0.001, d = 0.49) and second half (3.13 ± 0.99%, t0(71) = 2.89, p = 0.005, d = 0.37) of the experiment, with no difference between halves (t(71) = 1.05, p = 0.300, d = 0.12, BF_10_ = 0.219, Figure 2D). These results indicate that the TMS effects of LOC and OPA were robust over time.

Furthermore, to ensure that the results were not driven by specific features of one or a few objects or scenes, the LOC and OPA effects were compared between categories. There were no significant differences between the individual objects and scenes for the LOC effect (F(3,213) = 0.23, p = 0.859) or for the OPA effect (F(3,213) = 0.85, p = 0.465). Furthermore, all categories showed the effect in the expected direction (Figure 2E), although it did not reach significance for all individual categories tested separately (LOC effect: camera: p = 0.006, car: p = 0.084, chair: p = 0.048, shoes: p = 0.08; OPA effect: beach: p = 0.289, city: p = 0.064, forest: p = 0.008, kitchen: p = 0.006). This is likely a consequence of increased variability by selecting a subset of trials, which consequently reduces statistical power.

Finally, to exclude speed-accuracy trade-offs, we analyzed reaction time (RT) data. A 3×2 repeated measures ANOVA revealed no significant main effect of TMS site (F(2,142) = 0.839, p = 0.394) or category (F(1,71) = 3.28, p = 0.074), and no significant interaction of TMS site*category (F(2,142) = 0.11, p = 0.893). This indicates that RT did not differ between TMS conditions, neither for object nor scene stimuli, excluding speed-accuracy trade-offs.

## Discussion

The distinction between object and scene processing features prominently in cognitive neuroscience research (Oliva, 2013; Epstein, 2014). Indeed, neuroimaging studies have demonstrated that this distinction is one of the main organizing principles of human high-level visual cortex, with object-selective cortex representing object content (but not scene layout) and scene-selective regions representing scene layout (but not object content). Here, by testing a large sample (N=72) using an established paradigm (Dilks et al., 2013) and a pre-registered protocol, we provide conclusive TMS evidence supporting this distinction, with TMS over object-selective cortex (LOC) selectively interfering with object recognition and TMS over scene-selective cortex (OPA) selectively interfering with scene recognition. The results could not be explained by a speed-accuracy trade-off, and were consistent across time and individual object and scene stimuli.

What may explain the distinct functional roles of LOC and OPA, as revealed here? A likely possibility is that LOC contributed to object recognition by representing object shape or object contour, while OPA contributed to scene recognition by representing more globally distributed image properties such as surface texture (Cant and Goodale, 2007). Another possibility is that LOC and OPA differentially contributed to the tasks by processing different low-level visual input, such as spatial frequencies (Rajimehr et al., 2011), rectilinear features (Nasr et al., 2014), or retinal locations (Levy et al., 2001). Further research is needed to disentangle these contributions by using carefully controlled stimuli. For example, one fMRI study used a task manipulation to show increased activation in LOC when participants attended to the shape (flat vs convex) of a surface as compared to attending to its texture (rock vs wood; Cant and Goodale, 2011). The opposite result (texture>shape) was found in scene-selective cortex. It would be interesting to test whether LOC and OPA are causally involved in these tasks, for which visual input is equated.

While the current results provide strong evidence for distinct scene- and object-selective regions in visual cortex, this does not mean that scene- and object-selective pathways are functionally independent. In natural vision, scenes and objects are processed interactively, with scene context facilitating object recognition and object processing facilitating scene recognition. For example, when local object cues are degraded (e.g., a distant car in the mist), scene context can drive object recognition (Brandman and Peelen, 2017). Conversely, a clearly visible object can allow for recognizing an otherwise ambiguous scene (Brandman and Peelen, 2019). fMRI studies showed that degraded objects were more reliably represented in LOC when shown on a congruent scene background (Brandman and Peelen, 2017), and degraded scenes were more reliably represented in OPA when shown together with an intact object (Brandman and Peelen, 2019). In other words, object representations in LOC can be driven by scene cues, and scene representations in OPA can be driven by object cues. Future TMS studies could follow up on these results, testing whether LOC and OPA are causally involved in representing their preferred categories (objects or scenes) when these are merely inferred from non-preferred cues (scenes or objects).

To provide a near-exact replication of Dilks et al. (2013), we used the same behavioral task, stimuli, procedure, and TMS timing. However, one important difference between the studies is that we used group-average Talairach coordinates rather than individually-localized regions. This may implicate less spatial specificity in the present study, potentially decreasing the effect size (Sack et al., 2009). Nevertheless, the results of the two studies are remarkably similar (Figure 2B). In addition to the main results replicating, the effect sizes and variability are also quite comparable. The decrease in object recognition performance after LOC TMS was 4.7% (SD = 8.0%; Cohen’s d = 0.59) in the study by Dilks et al. and 3.9% (SD = 8.3%; Cohen’s d = 0.47) in the present study. The decrease in scene recognition performance after OPA TMS was 4.9% (SD = 5.7%; Cohen’s d = 0.86) in the study by Dilks et al. and 3.8% (SD = 6.4%; Cohen’s d = 0.59) in the present study. These results show that LOC and OPA can be targeted reliably based on group-average fMRI coordinates, providing an easy and cost-effective way to investigate the LOC and OPA when an fMRI scan is not feasible.

Finally, TMS timing in the current study was rather aspecific with a window of 500 ms and a frequency of 10 Hz. Several studies have suggested that the LOC represents object category between 120 and 200 ms after stimulus onset (Koivisto et al., 2011; Isik et al., 2014; Carlson et al., 2013; Cichy et al., 2014; Kaiser et al., 2016). Similarly, the OPA represents scene layout around the same time window (Cichy et al., 2017; Henriksson et al., 2019). Based on these values, TMS timings could be optimized to disrupt object and scene recognition more powerfully, with a narrower window and a higher frequency, for example TMS between 100 and 300 ms after stimulus onset with a frequency of 20 Hz. Additionally, single- or double-pulse TMS at specific latencies should be used to test the time points at which these regions are causally involved in the recognition of objects and scenes (Koivisto et al., 2011; Pitcher et al., 2012; Reeder et al., 2015).

In summary, the current study provides causal evidence for a double dissociation between object-selective LOC and scene-selective OPA, successfully replicating Dilks et al. (2013). By showing that these regions can be reliably targeted with TMS, even when using group-average fMRI coordinates, we can now address new questions regarding the functionality of these regions. One interesting avenue for future research is to move beyond isolated object and scene processing and test the causal time course of LOC and OPA during naturalistic situations, in which scene and object processing interact and mutually inform each other.

## Acknowledgements

This project has received funding from the European Research Council (ERC) under the European Union’s Horizon 2020 research and innovation programme (grant agreement No 725970). We would like to thank Daniel Dilks for providing the stimuli and task.

